# Transcriptomic analysis reveals lipid metabolism and macrophage involvement associated with nintedanib treatment in a rat bleomycin model

**DOI:** 10.1101/2024.10.30.620876

**Authors:** Martina Bonatti, Vanessa Pitozzi, Paola Caruso, Silvia Pontis, Maria Gloria Pittelli, Caterina Frati, Denise Madeddu, Federico Quaini, Costanza Anna Maria Lagrasta, Ilaria Minato, Emanuela Bocchi, Maurizio Civelli, Gino Villetti, Marcello Trevisani, Barbara Montanini

## Abstract

**INTRODUCTION:** Idiopathic pulmonary fibrosis (IPF) is a progressive and irreversible lung disease with a poor prognosis. While pirfenidone and nintedanib offer some benefits, they cannot cure IPF. Nintedanib inhibits various proliferative pathways and has antifibrotic effects, but its molecular mechanisms and impact on the lung transcriptome in vivo remain unclear. This study aims to evaluate nintedanib’s transcriptomic profile in a rat model of bleomycin-induced lung fibrosis.

**METHODOLOGY/PRINCIPAL FINDINGS:** Lung fibrosis was induced by two intratracheal administrations of bleomycin. Nintedanib protocol included three weeks of daily oral treatments beginning seven days after the first bleomycin dose. Left lungs were processed for histological evaluation using an automated fibrosis quantification system and the Ashcroft Score, while the right lungs were used for RNA sequencing to conduct differential expression and correlation network analysis (WGCNA). WGCNA modules were examined by cell and pathway enrichment analysis. Lipid peroxidation was assessed through the measurement of malondialdehyde in right lung lysates.

Bleomycin induced significant fibrotic lesions, as confirmed by the histological evaluations. Nintedanib reduced fibrotic lesion size by about 15% and decreased severe Ashcroft scores. When compared to controls, the number of differentially expressed genes decreased from over 2000 to barely more than 400 after nintedanib treatment. WGCNA identified two gene clusters correlated to histological parameters, with nintedanib-treated animals showing gene expression levels similar to control animals. One cluster was associated with mesenchymal cells and extracellular matrix-related pathways, in line with the known anti-fibrotic effect of nintedanib. The second cluster, involving principally macrophages, was related to lipid metabolism, potentially uncovering a new mechanistic role of nintedanib in modulating lung fibrosis.

**CONCLUSIONS/SIGNIFICANCE:** The mechanisms involving macrophages and lipid metabolism, influenced by nintedanib in this study, may open new research directions to better inquire the role of this cellular type in tissue repair and pathological lung fibrosis.

## INTRODUCTION

Idiopathic Pulmonary Fibrosis (IPF) is a deadly Interstitial Lung Disease (ILD) with unclear pathogenic clues, characterised by damaged lung parenchyma architecture, progressive restrictive-ventilatory limitation, hypoxia, dyspnoea, and cough [1]. Its development is believed to be influenced by both genetic and environmental factors and leads to its distinctive histopathologic and radiologic pattern defined Usual Interstitial Pneumonia (UIP) [2]. Only a few pharmacological options are available, pirfenidone and nintedanib, and there is no long-term effective treatment or curative medicine for IPF, highlighting an enormous medical need [1]. Nintedanib is a small molecule that acts at the intracellular level and inhibits a variety of multiple tyrosine kinase receptors, with the following being the most significantly impacted: vascular endothelial growth factor receptor (VEGFR), platelet-derived growth factor receptor (PDGFR), and fibroblast growth factor receptor (FGFR) [3]. Nintedanib gained approval for treating IPF thanks to its efficacy in reducing acute exacerbations, minimizing the risk of disease progression, and slowing the decline of Forced Vital Capacity (FVC) in patients with lung fibrosis [1, 4]. Like other tyrosine kinase inhibitors, it plays a regulatory role in essential processes such as differentiation, growth, metabolism, and apoptosis [5].

Animal models that resemble pulmonary fibrosis could contribute to the discovery of novel potential therapeutic approaches for complex lung diseases, even though there is no adequate and relevant preclinical model recapitulating IPF [6]. However, it is widely accepted and pointed out in guidelines that the bleomycin (blm) murine model stands out as the best-characterized and extensively studied IPF animal model for preclinical testing of active compounds on IPF [6, 7]. Transcriptome profiling and advanced molecular biology techniques, such as single-cell omics, offered critical insights into cellular dysfunctions in complex pathologies, and recent successes in applying these methods to lung fibrosis models and IPF-related experiments provided valuable translational perspectives [8–12].

In IPF and in fibrotic models, nintedanib is reported to interfere with a number of biological processes critical for the disease’s development and progression, including the fibroblast-to-myofibroblast transition (FMT) and the excessive production and deposition of extracellular matrix (ECM) components such as collagen [1]. Particularly, its effect on fibroblasts is widely documented [13] and Shue et al. reported a well-defined set of significantly dysregulated genes associated with nintedanib treatment in IPF fibroblasts [10]. Nevertheless, the understanding of the full spectrum of nintedanib’s effects remains incomplete. It is acknowledged that nonreceptor tyrosine kinases, as well as unidentified targets and cellular processes, may play pivotal roles in mediating antifibrotic effects *in vivo* [13, 14]. To our knowledge, this is the first comprehensive transcriptome and histomorphometric analysis of a rat double-hit blm model of lung fibrosis treated with nintedanib, with the aim to uncover the mechanisms underlying nintedanib benefits and potentially identify new targetable therapeutic pathways.

## RESULTS

### Therapeutic effect of nintedanib on blm-induced lung fibrosis

At day 28, blm produced a lung fibrosis consisting of dense collagen agglomerates that replaced normal lung parenchyma, making the alveoli difficult to recognize. Nintedanib treatment appeared to reduce the density of fibrotic area and collagen deposition with alveoli still easy to recognize (Fig. 1B). Automated quantification of lung fibrosis showed fibrotic tissue covered around 25% of the total lung area in the BLM group, while nintedanib-treated animals had only 15% lung damage. (Fig. 1C). The Ashcroft score analysis revealed a median 19% reduction compared to the BLM group (p ≤ 0.01, Fig. 1D). Moreover, the frequency of moderate (4-5) scores dropped in favour of mild ones (2-3) in the NINT group. As expected, the BLM group showed high variability, with certain animals displaying more severe and widespread lesions than others. Nevertheless, despite these notable disparities, the variability did not impact the statistical significance of histologic parameters documented in BLM group compared to SAL. The mean values of these parameters were comparable to other studies on a similar rat model.

**Fig. 1.**
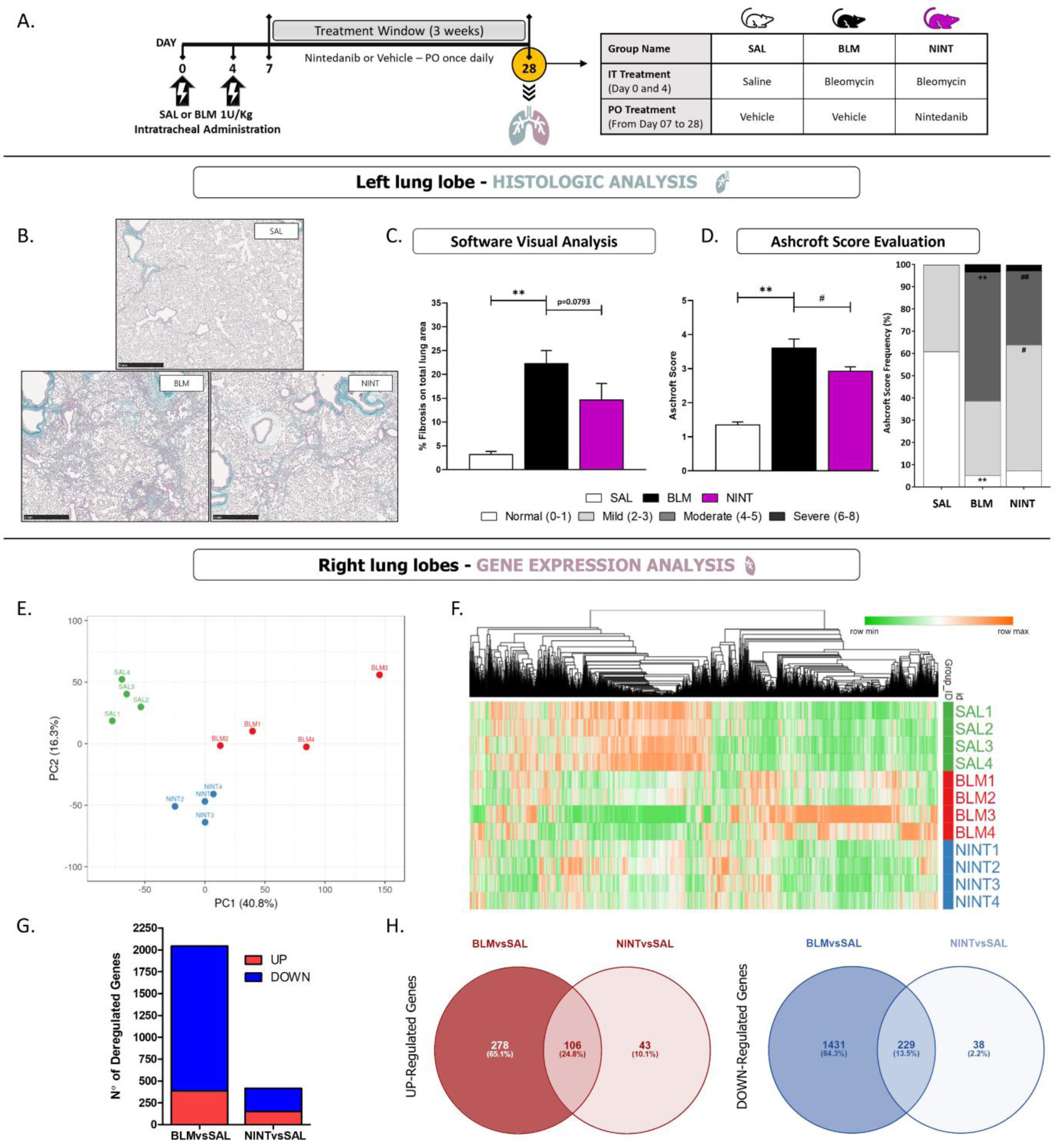
Comprehensive overview of the rat model. (A) Graphical description of the rat model and the experimental set-up, indicating the workflow and timeline of the different interventions. (B) Representative Masson’s trichrome-stained lung tissue sections at the end of the experiment, 28 days after the first BLM instillation. (C) Quantification of pulmonary fibrosis using the automated software visual analysis and (D) Ashcroft score evaluation, expressed as mean values or percentages of the frequency across different score ranges. **p.val ≤ 0.01 vs. SAL, ^##^p.val ≤ 0.01 vs. BLM, and ^#^p.val ≤ 0.05 vs. BLM. The histological evaluations were performed on all the animals in the experiment (n = 8). (E) Graphical representation of the sample variably using the PCA. The normalised counts of all genes were utilised for this purpose. (F) The heatmap displays the normalised counts of all animals that performed transcriptome analysis. Only genes that exhibited a statistically significant de-regulation in at least one of the comparisons were chosen for the heatmap. The columns were hierarchically clustered using a Euclidean distance metric. (G) Bar graphs depicting the number of genes that were either UP-regulated or DOWN-regulated in the two groups, BLM and NINT, in comparison to the control group (SAL). (H) Venn-diagrams of the differential expressed genes among the different groups.

### Blm induced the production of lipid-laden AMs

The majority of the Masson’s trichrome stained sections from the animals treated with blm exhibited clusters of larger and atypical AMs with a foamy appearance resembling lipid-laden AMs, prevalently localized in areas of thickened tissue. These structures are absent in the severely injured fibrotic regions where the alveoli are no longer visible. A detail of these histologic findings is reported in Fig. 4A. These structures are not present in any of the control animals.

### Overall impact of nintedanib on blm-altered lung transcriptome

The Principal Component Analysis (PCA) performed on the RNAseq data revealed a clear separation among the analysed groups (Fig. 1E). The animals in the NINT group (light blue), like the controls (green), appeared to be tightly grouped together. The four animals in the BLM group (red) were more dispersed on the PC1.

This variability among BLM group members can also be clearly visualized in the transcriptome profiles (Fig. 1F). In detail, BLM1 and BLM2 show relatively moderate alterations compared to the control group, while BLM3 and BLM4 showed a more altered profile compared to the controls.

The number of DEGs ranged from 416 (NINT vs SAL) to 2044 (BLM vs SAL) (Fig. 1G). Down-regulated genes significantly outweigh up-regulated genes, as previously observed for this model at this time point [8]. The Venn diagrams presented in Fig. 1H show the overlap of up-regulated (in red) and down-regulated (in blue) genes between the two analyzed comparisons. The 106 up- and 229 down-regulated genes in common reveal the transcriptome segment impacted by the blm but unaffected by the therapeutic effect of nintedanib. 278 up-regulated and 1431 down-regulated genes are significantly modified by blm only. This finding highlights a deregulated portion of the blm-altered lung transcriptome that nintedanib treatment may have restored. However, 43 up-regulated and 38 down-regulated genes underscores the existence of nintedanib-sensitive genes independently of the blm effect.

### Identification of specific gene expression signatures associated with nintedanib effect

We employed a network-based approach to identify specific transcriptional profiles. The signed WGCNA identified 14 modules of co-regulated genes. All the modules with their correspondent characteristics are graphically reported in Fig. 2. The first four detected modules accounted for 81.25% of all genes, with Module 1 alone accounting for roughly half of them, while the next ten modules included less than 6% each.

**Fig. 2.**
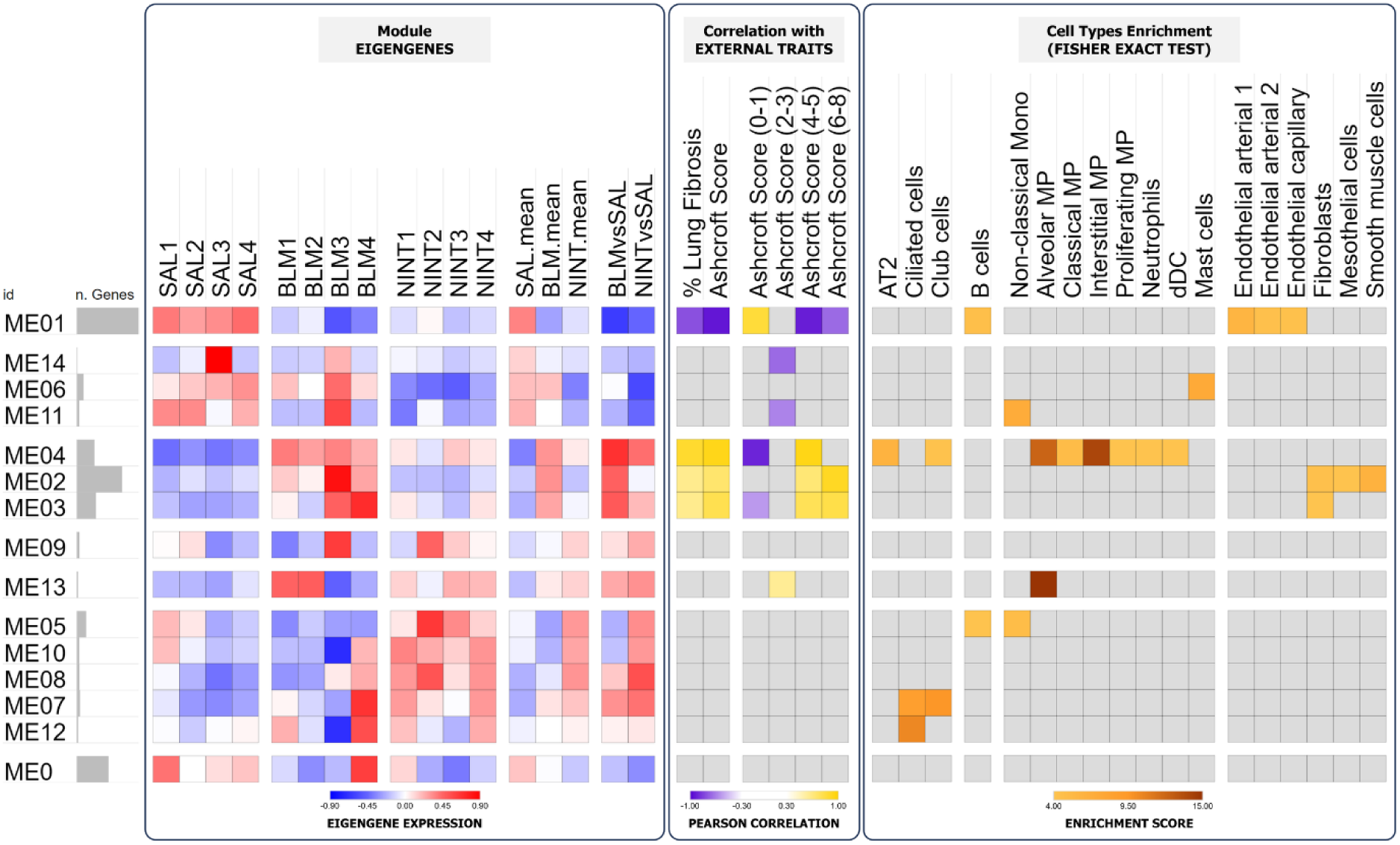
Gene modules identified using signed WGCNA. Representative heatmaps of each WGCNA module. The first heatmap (Module EIGENGENES) is coloured based on the module eigengenes obtained for each animal. Additionally, the mean of each group and the ratio of the mean values for the BLM and NINT groups against the control group (BLM-NINT vs. SAL) are reported. The second heatmap (Correlation with EXTERNAL TRAITS) depicts Pearson correlations obtained during WGCNA analysis with the indicated external traits. The third heatmap (Cell Types Enrichment) displays the enrichment of cell-type-specific gene lists for genes with module membership ≥ 0.8 (cell-types are grouped based on the lineage, i.e. epithelial, lymphoid, myeloid and stromal). In these last two heatmaps, non-statistically significant values are represented in grey.

Module eigengenes (MEs) are representative of the expression profile of co-regulated genes within each module. ME01 showed elevated expression in the control group, while the BLM and NINT groups displayed reduced expression levels. However, the BLM group showed variability in the extent of downregulation observed. Many of the significantly down-regulated genes from the DEG analysis fell into Module 1. ME02 exhibited increased expression levels in animals of the BLM group compared to controls, with notable variation, particularly underscored by very high eigengene expression for BLM3. The levels of expression of the NINT animals were comparable to that of the SAL group. In ME03, a similar pattern emerged, although NINT animals appeared closer to the expression levels of BLM1 and BLM2 than the SAL group. In contrast to other modules, ME04 stood out as unique. Notably, it is the only module where the expression levels were uniform across the entire BLM group and higher than those in the SAL group. Furthermore, in this ME, the NINT group showcased an intermediate expression profile, positioned between the SAL and the BLM groups.

The Pearson correlation of the first four MEs significantly correlated in varied intensities and orientations with the values of the Ashcroft Score and percentage of fibrotic tissue evaluated on lung sections of the same animals (Fig. 2). The correlations with the varying degrees of lung fibrosis severity revealed differences in correlation patterns, with ME02 and ME03 more closely related to severe grades of fibrosis.

The enrichment of cell-type-specific signatures within modules unveiled the specific cell types associated with the main modules of interest (Fig. 2). Module 1 appeared to be characterized by genes belonging to the signatures of vascular endothelial and B cells (score = 4.93 and 4.68, respectively), while Module 2 was distinguished by genes associated with cell types crucial for the fibrotic process, such as smooth muscle cells, fibroblasts and mesothelial cells (score = 6.31, 4.45 and 4.14, respectively). Module 3 also showed a slight association with fibroblast signature (score = 4.05). Module 4 exhibited highly statistically significant scores associated with various types of resident macrophages (score = 12.36 for AMs and score = 14 for interstitial macrophages) and alveolar type 2 cells (AT2; score = 6.14), as well as other immune-inflammatory cells (dendritic cells and neutrophils, scores = 4.16 and 4.12, respectively).

### Pathway analysis of the WGCNA modules

The pathway analysis revealed numerous signalling pathways, and several biological processes enriched with high statistical significance (Fig. 3), the complete list is reported in supplementary material (Table S1). Module 1 was enriched in pathways related to gene regulatory processes, with very high statistically significant enrichment scores (q-val. < 10^-30^) for terms such as chromatin organisation, mRNA metabolic processes, histone binding, and transcription coregulator activity, with the highest enrichment for chromatin activities (q-val. < 10^-61^). This module reached very high statistical significance (q-val. < 10^-25^) also for terms related to cytoskeletal dynamics, such as cell projection assembly, microtubule cytoskeleton organisation, and the RHO GTPases cycle. Furthermore, several interesting terms involved in cellular and genomic stability were found to be enriched, like DNA damage response and ATP-dependent activity.

**Fig. 3.**
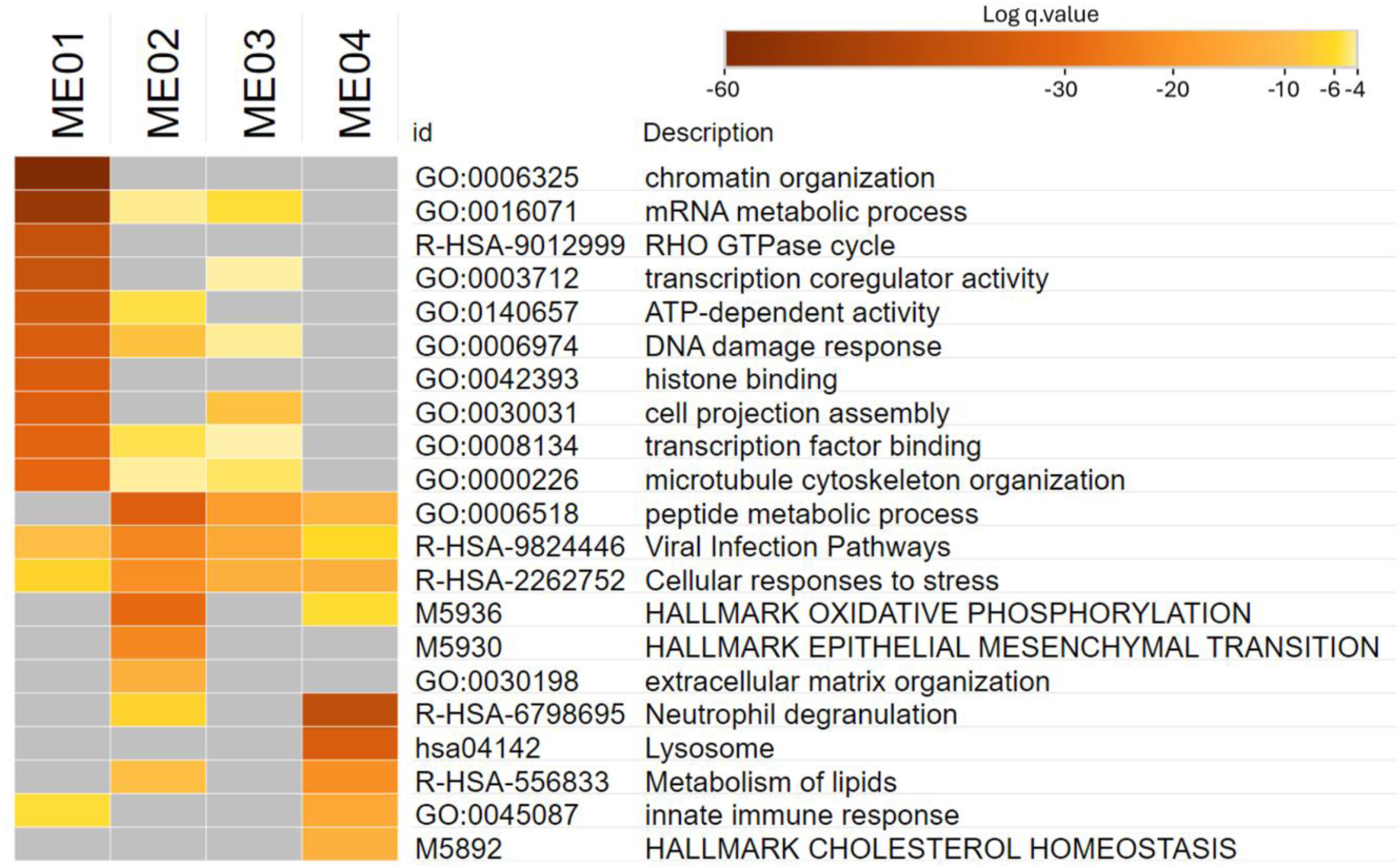
Pathway enrichment analysis of WGCNA modules. The heatmap displays Log q values derived from pathway analysis conducted using Metascape. The most noteworthy pathways and terms were selected from the original comprehensive list, starting with the most significant ones. These selections are either uniquely associated with a specific module or commonly enriched among a group of modules.

Modules 2, 3, and 4 were notably more similar to each other in terms of enriched pathways. These common pathways highlighted processes and mechanisms related to cellular and viral translational control, ranging from ribosomal subunit formation and activity to protein synthesis and modification processes, including peptide metabolic and biosynthetic processes, translation, and amide biosynthesis. Some of these terms were also enriched in Module 1. The common line of Modules 2, 3, and 4 focused on events related to translation and protein synthesis, whereas the one shared with Module 1 delved into the dominance of mRNA metabolism, splicing, and associated processes critical for gene expression and cellular regulation.

In addition to these common pathways, Modules 2 and 4 had uniquely enriched terms. Module 2 was enriched for pathways related to cellular respiration and energy metabolism, such as oxidative phosphorylation and respiratory electron transport of mitochondria. Moreover, there was a strong enrichment for pathways linked to the fibrotic process, like epithelial-mesenchymal transition and extracellular matrix organization. Module 4 was enriched in mechanisms related to the innate immune response, with very high statistical significance (q-val. < 10^-30^) for neutrophil degranulation and lysosomal activity, and pathways governing lipid metabolism, including cholesterol homeostasis and lipid biosynthesis, alongside processes like monocarboxylic acid metabolism and adipogenesis. The results were summarized in the graphical abstract, which was partially generated using BioRender.com.

### Comparison with other relevant studies

A gene set overrepresentation analysis provided statistically significant enrichment scores for Modules 1, 2, and 4 when compared to our previous time course study [8], were we investigated the temporal fluctuations in gene expression during the progression of lung fibrosis in the same rat model. Specifically, Module 1 is enriched in genes belonging to the late response, while Modules 2 and 4 are enriched in genes belonging to the long-lasting one. A complete list is reported in supplementary materials (Table S2).

With the GSEA, we obtained a statistically significant enrichment score for an *in vitro* experiment published by Sheu et al. [10], where gene expression changes associated with nintedanib treatment in IPF fibroblasts were assessed. Specifically, the statistical significance was achieved by comparing Module 2 with the list of down-regulated genes after nintedanib treatment. The associated GSEA leading-edge analysis provided a list of 40 genes for this enrichment. All the results of the GSEA and leading-edge analysis are reported in supplementary materials (Table S3-4).

The gene set overrepresentation analysis, employing cell-type-specific markers from a IPF single-cell RNA-sequencing study [11], revealed robust and statistically significant enrichment scores for Modules 2 and 4 for some specific IPF-related cell types. Specifically, Module 2 exhibited enrichment for genes associated with fibroblasts and proliferating cells, while Module 4 demonstrated significance for specific signatures of AT1/club, AT2, low-quality basal, and two distinct types of macrophages (SPP1hi and FABP4hi, with 18.91 and 8.62 as enrichment score respectively. We also obtained other statistically significant enrichments for minor modules; the complete list, along with all the enrichment scores and relevant statistics, is reported in the supplementary material (Table S5).

### Nintendanib attenuates blm-induced lipid peroxidation

Considering the involvement of lipid-related pathways, in addition to known oxidative state-inducing mechanisms (e.g., mitochondrial respiration and inflammation) and the reported presence of lipid-laden AMs in blm-treated lungs (Fig. 4A), we explored the potential implications of lipid peroxidation in our model, as well as its intersection with nintedanib. To do this we analyzed the concentration of MDA, as it represents one of the most frequently utilized approach to quantify lipid peroxidation and a prominent indicator of oxidative damage. As reported in Fig. 4B, a statistically significant increased MDA concentration was observed in BLM group compared to SAL. The NINT group also displayed an increased MDA concentration compared to SAL, although to a lesser extent compared to BLM, indicating that nintedanib partially attenuates lipid peroxidation and consequently oxidative damage.

**Fig. 4.**
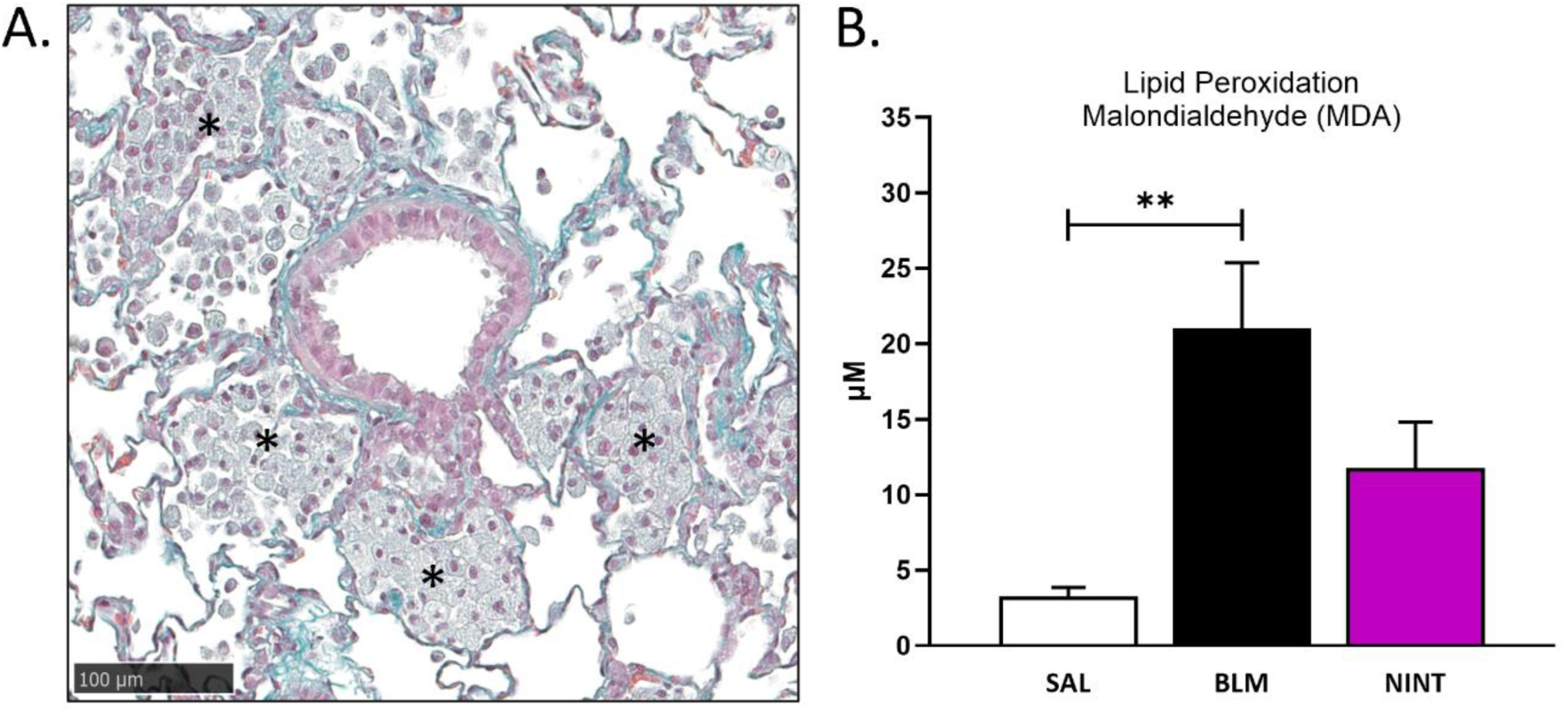
Effect of nintedanib on lipid associated lung damage induced by blm. (A) Clusters of lipid-laden alveolar macrophages (* foam cells) are illustrated on Masson’s trichrome-stained lung tissue sections from blm-treated rats (BLM group). Original magnification X20. (B) Bar graphs showing malondialdehyde (MDA) concentration in lung homogenates from all the groups. Data reported as mean ±SEM, **p.val ≤ 0.01 vs. SAL.

## DISCUSSION

Despite substantial advances in understanding the intricate molecular features of chronic lung diseases and the use of advanced multi-molecular techniques, progress in identifying effective treatments for IPF has been unsatisfactory. Promising candidates have often failed in clinical trials, highlighting the urgent need for better therapies. Diverse animal models, ranging from pharmacologically induced to genetically engineered mice, have provided valuable insights into the biological mechanisms of IPF [7, 15–20], even though there is currently no *in vivo* model that can accurately mimic the complexity of human IPF. Despite the limitations of various animal models, the blm model revealed the beneficial effects of nintedanib, which successfully crossed all clinical stages until approval [21], demonstrating efficacy in retarding disease progression in IPF patients [22]. The main molecular targets of nintedanib, as well as the core part of its mechanism of action, are well-documented [13], and many published studies determined the effects of nintedanib on lung fibroblasts, which are key effector cells in the pathogenesis of fibrosis [10, 23, 24]. However, thanks to recent single-cell approaches [11, 12], we now know that other important cellular players are involved in IPF. Understanding all the biomolecular details of the antifibrotic mechanisms of nintedanib could be pivotal for advancing future therapeutic approaches; however, an in-depth biomolecular analysis of its effects on lung fibrosis *in vivo* is lacking. Therefore, to delve into the full spectrum of molecular mechanisms associated with nintedanib treatment in IPF, we employed a transcriptomic approach to analyze mRNA alterations in a blm-induced lung fibrosis rat model following a therapeutic regimen of nintedanib. A similar approach was performed by Sheu et al. [10], that evaluated changes in mRNA and miRNA profiles on IPF fibroblasts exposed to two doses of nintedanib. However, the proposed system is monocellular, which limits the information on the molecular interplay between different cell types and cell-matrix communication. In our study, many DEGs were identified as being associated with nintedanib treatment, along with specific molecular signatures accounting for the beneficial effects of the drug. These findings may provide valuable insights to identify new potential therapeutic targets of IPF that can be selectively addressed to develop more tolerated drugs.

Overall, our research revealed significant variations in the impact of blm on lung injury severity. This variability was evidenced by histological parameters and by the transcriptomic profiles. Indeed, BLM1 and BLM2 showed moderate alterations, with expression profiles that were not completely distinct from those of the control group. Conversely, BLM3 and BLM4 exhibited more severely damaged lungs, and a substantially altered transcriptomic profile. The variability in fibrosis severity induced by blm in rats has been already reported and extensively examined by Persson et al. [25]. They found two animal subgroups, low and high responders, depending on their reactions to blm. Both showed inflammation, but low responders demonstrated early healing, whereas high responders increased lung and lesion volume. However, despite the variability observed in our study, we were able to identify specific gene expression signatures that reflected the effects of nintedanib. The ME4 exhibited a consistent pattern of increased gene expression in animals treated with blm compared to the control group, and unlike all the other module eigengenes, low variability in gene expression levels was documented among BLM animals. For this ME, the gene expression levels of the NINT group were located halfway between those of the BLM and SAL groups, indicating a mild yet discernible impact of nintedanib. Moreover, this group of genes were not influenced by the varying levels of severity to blm. The ME2 also provides valuable insights into nintedanib’s effects. Despite the varying gene expression levels within the BLM group, with significantly higher expression observed in the more severely injured BLM3 and BLM4 animals, all members of the NINT group exhibited similar levels of gene expression as the control animals. This suggests that nintedanib effectively normalizes gene expression levels regardless of the initial severity of lung injury, indicating a strong molecular interference on genes of Module 2. In contrast, in ME1, the animals in the BLM group that have a less severe lung injury profile (BLM1 and BLM2) exhibit gene expression levels comparable to those of all the animals treated with nintedanib. Therefore, conclusive evidence regarding the efficacy of nintedanib on these genes cannot be provided in the absence of knowledge about the initial severity of NINT animals.

Modules 2 and 4 emphasize two distinct beneficial effects of nintedanib, which are graphically summarized in Fig. 5. The effect highlighted by Module 2 is arguably the most anticipated one. Indeed, Module 2 is enriched in genes associated with cell types that are typically altered and pathogenically involved in IPF, including fibroblasts, mesothelial and smooth muscle cells, which are generally targeted by anti-fibrotic drugs. Moreover, this module is enriched in genes reported to be modified by nintedanib in the fibroblast model of Sheu et al. [10]. Not surprisingly, this module features pathways related to extracellular matrix organization and epithelial to mesenchymal transition, processes that have been reported to be affected by nintedanib in a fibrotic context [13]. Regarding other pathways and processes that are enriched in this module, especially those related to mitochondrial involvement and oxidative phosphorylation, there is also evidence indicating their contribution to lung fibrosis [26–28]. However, the effect of nintedanib on these processes remains to be elucidated.

**Fig. 5.**
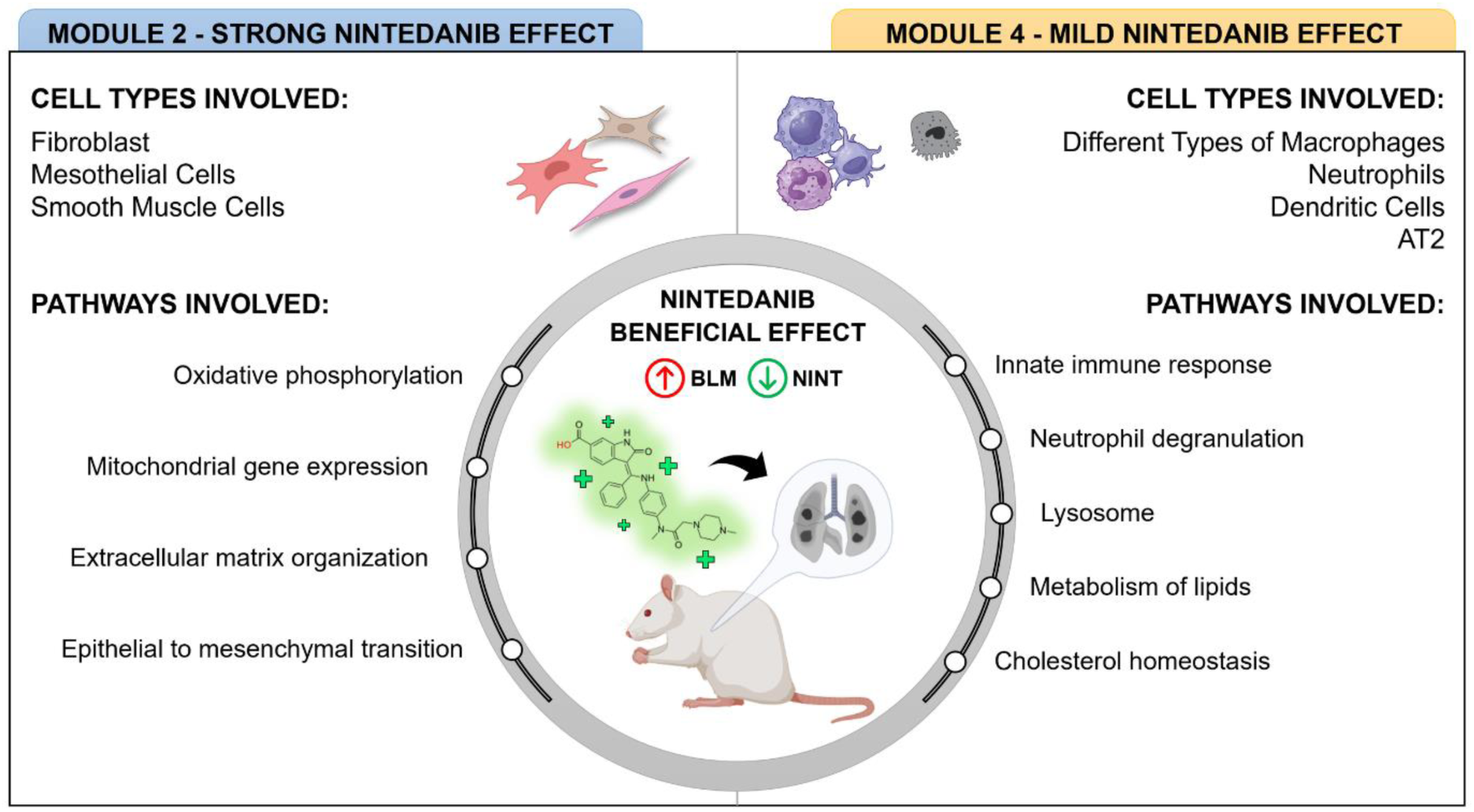
Summary of the molecular effect of nintedanib on the reported rat model. Schematic representation partially created with BioRender.com.

Module 4 is significantly enriched in AMs and AT2 cells associated gene signatures and shows enrichment in pathways related with lipid metabolism and cholesterol homeostasis. These findings collectively hint at a possible dysregulation of lipid-related pathways, particularly during the interaction between AMs and AT2 cells [29]. Moreover, histological features of our blm-treated rats included the presence of agglomerates of lipid-laden AMs (Fig. 4A). These cells have been reported to be extremely relevant in different contexts of human lung fibrosis [30–32], as well as in blm-induced lung fibrosis models [29, 33, 34]. These findings imply that there is a mechanism primarily involving AMs and lipids that contributes to the fibrotic process generated by blm. This mechanism is independent of lesion severity and is affected by nintedanib. Increasing evidence suggests that pulmonary fibrosis is caused by lipid metabolism issues, such as anomalies in low-density lipoprotein (LDL) metabolism and alterations in the oxidative status of various lipids [29, 32, 35]. Furthermore, there is a notable association and a commonality of lipid risk factors between pulmonary fibrosis and disorders characterized by substantial abnormalities in lipid metabolism, such as atherosclerosis and obesity [35]. Lipid-laden AMs, also known as “foam cells”, are established pathogenic features and have been shown to be extremely relevant in animal models of lung fibrosis [29, 32–34, 36], as well as histological specimens from humans with chronic fibrotic lung disorders, such as IPF [29–31]. In addition, other significant lipid-related cell types (such as AT2s) and other highly specific IPF-related macrophages are being investigated as potential initiators and contributors to the pathogenic mechanism of lung fibrosis [11, 12, 35]. The combination of these actors and aspects resembles a pathological mechanism affecting macrophages in another disease, atherosclerosis [37]. In the context of lung fibrosis this mechanism has been elucidated by Romero and colleagues under the name of pneumocyte–macrophage paracrine lipid axis [29]. Data of the present investigation strongly indicate the involvement of this axis in our model, potentially underlying the positive effect of nintedanib. According to this view, lipids produced by AT2 cells to generate the surfactant are oxidized in the oxidant environment created by the pathologic conditions. Our data highlighted a dysregulation of various oxidative stress-related pathways, including inflammation and oxidative phosphorylation, and elevated MDA levels in blm-treated rats. The high concentration of oxidized lipids alters macrophage activities, causing up-regulation of lipid receptors and subsequent lipid internalization and accumulation. Macrophage scavenger receptors, such as Orl1 (oxidized low density lipoprotein receptor 1) and Scarb2 (scavenger receptor class B, member 2 or CD36 Antigen-Like 2), recognize and internalize oxidized lipids [37]. The genes coding for these two receptors both belong to module 4 in our model, and resulted statistically different in BLM vs SAL, with log_2_FC values of 1.27 and 0.92, respectively. The same observations were reported in a model of silica dust, where the authors demonstrated elevated levels of intracellular lipids, specifically oxidized LDL (ox-LDL), and high transcript levels for the lipid scavenger receptor [32]. The accumulation of oxidized lipids in the extracellular space is sufficient to trigger the formation of lipid-filled macrophages [29], such as those we reported in our histological images of blm-treated animals. Once internalized by macrophages, oxidized lipids (such as ox-LDL) can accumulate in the lysosome to be converted into free-cholesterol, as indicated by the ultrastructural research conducted by Bobryshev and colleagues [37]. Lysosome is the second most significant enriched term (q.val = 10^-34^) observed here among Module 4 genes, where it is uniquely enriched. Once taken up by macrophages, cholesterol and other lipids have been shown to activate multiple pro-fibrotic pathways and particularly to induce TGF-β1 signalling [29].

As mentioned earlier, in Module 4, nintedanib exhibits a mild, yet beneficial effect. Thus, in the light of the emerging axis of lipid metabolism driving pulmonary fibrosis, nintedanib could exert a protective effect on lipid oxidation, supported by already published data. Indeed, Li and colleagues [38] demonstrated that two progressive doses of nintedanib can reverse the increased production of total cholesterol, free cholesterol, and pro-inflammatory cytokines that were induced in ox-LDL-treated human umbilical vein endothelial cells (HUVECs). Moreover, the increase in reactive oxygen species and MDA, as well as the declining activity of glutathione peroxidase in ox-LDL-treated HUVECs, were significantly abolished by the two nintedanib concentrations [38]. We observed a similar degree of reduction of MDA in lung homogenates from nintedanib-treated animals compared with the BLM group. The assertion that nintedanib positively affects pathways related to lysosome and lipid metabolism is supported also by a separate study that evaluated the effects of nintedanib on gastric cancer cells [39]. Indeed, examination of nintedanib-treated cells by tandem mass tags proteomics revealed the presence of more than 800 differently expressed proteins, which were enriched in multiple pathways, including fatty acid metabolism and lysosome [39].

The findings and references pertain more to the context of general pulmonary fibrosis and related models. However, the importance of different types of macrophages is gaining increasing attention in the specific IPF context. In more detail, Morse et al. [11] emphasize in their publication that two macrophage populations are present in diseased areas of the IPF lungs, designated as SPP1^hi^ and FAPB4^hi^. The first population is part of the interstitial macrophage compartment, while the second makes up most of the macrophages in alveoli [11]. SPP1 (secreted phosphoprotein 1), which is known to support macrophage proliferation, was observed to be deposited in a striking manner in fibrotic IPF lower lobes, where it was associated with fibroblastic foci [11]. FABP4 (Fatty Acid-Binding Protein 4) encodes for a fatty acid-binding protein, and its roles include fatty acid uptake, transport, and metabolism. When expressed by macrophages, it induces inflammation associated with obesity and atherosclerosis [40]. The gene markers of both macrophage populations are enriched in Module 4 and are up-regulated in our model (BLMvsSAL log_2_FC = 3.79 Spp1 and 1.23 FABP4). Interestingly, the FAPB4^hi^ macrophages are linked to lipid metabolism, and we speculate their potential to develop in foamy macrophages, perhaps representing an early/intermediate state of diseased macrophages. Indeed, Morse et al. report that despite increased proliferation, the overall percentage of FABP4^hi^ macrophages decreases in the most diseased IPF lower lobes, suggesting these cells either die or transition to SPP1^hi^ macrophages [11]. This is highly consistent with our model, as foamy macrophages are mainly identified in pseudo-isolated areas, far from the most damaged areas filled with scar tissue and collagen deposition. Furthermore, after nintedanib treatment, SPP1 is reduced by half (NINTvsSAL FC = 2.09), whereas FABP4 increases slightly (NINTvsSAL FC = 1.66), implying that the impact of nintedanib may work to reduce the development of SPP1^hi^ macrophages.

In conclusion, in our rat model of blm-induced lung fibrosis, we demonstrated the therapeutic and molecular impact of nintedanib using a combined histological and transcriptomic approach. This is the first time, to our knowledge, that a transcriptomic analysis on the effect of nintedanib has been conducted in the rat blm model as a surrogate for human lung fibrosis. In addition, using a network-based cluster analysis, we successfully identified two distinct gene expression patterns underpinning two diverse molecular effects of nintedanib on lung fibrosis. One involves mesenchymal cells, influencing pathways related to extracellular matrix organization, epithelial-to-mesenchymal transition, oxidative phosphorylation and mitochondria. The other, with a more modest impact, primarily engages macrophages and AT2 cells, and affects lipid metabolism and oxidation. To our knowledge, this is the first report of the mechanistic effect of nintedanib on lipid metabolism in the context of pulmonary fibrosis.

We acknowledge that our study has some limitations due to the small number of animals employed and the variability of the blm effect. In addition, we recognize that the observed impact of nintedanib on lipid metabolism and oxidation seems minor in comparison to the effect on fibroblasts and other well-known fibrosis-related pathways. However, the variability in the effect of blm was critical in identifying new prospective paths of nintedanib activity independently from blm severity. If the model had a uniform response, this small influence would have been statistically overshadowed by the other massive effect displayed by nintedanib on our blm model.

The specific roles and molecular mechanisms of lipid metabolism reprogramming in pulmonary fibrosis and its interference by nintedanib remain inadequately dissected. However, given the beneficial effects of this molecule in IPF patients and on lung fibrosis in general, as well as the growing importance of lipid metabolism in fibrotic lung diseases, we strongly believe that a thorough investigation of this topic is needed to uncover new therapeutic strategies specifically targeting the lipid machinery.

## MATERIALS AND METHODS

### Experimental Design

All the procedures involving animals were reviewed, approved, and authorized by 1) The Italian Public Health System for Animal Health and Food Safety within the Italian Ministry of Health (authorization number 246/2021-PR) and by 2) the Chiesi Body for the Protection of Animals (OPBA) committee. Experiments were performed in full compliance with the European ethics standards in conformity with directive 2010/63/EU, Italian D. Lgs 26/2014, the revised “Guide for the Care and Use of Laboratory Animals” (Guide for the Care and Use of Laboratory Animals, 1996).

Male Sprague-Dawley rats (Charles River, Italy) were intratracheally injected with blm (1 U/kg/0.5 ml; bleomycin sulfate; Baxter Oncology GmbH, Germany) at day 0 and after 4 days using a Penn Century Microsprayer (Penn-Century Inc., Philadelphia, USA) to induce pulmonary fibrosis. The same procedure was performed on control animals using a saline solution. More information regarding the model was published before in Bonatti et al. (2023) [8]. Seven days after the first blm dose, the animals were randomly divided into two groups, one of which received nintedanib 100 mg/kg (nintedanib free base – Carbosynth, Compton – UK; Group name: NINT) and the other received its vehicle (Methylcellulose 0.5% in water; Group name: BLM). Nintedanib and the vehicle were administered daily by oral gavage (per os, PO) for three weeks. During the weeks of treatment, the animals treated with saline received the vehicle only (Group name: SAL). A graphical representation of the experimental design and of the different groups is reported in Fig. 1A. Each of the three groups contained six animals. All the animals were anaesthetized with thiopental (200 mg/kg/10 ml; pentothal sodium; MSD Animal Health Srl) and sacrificed by bleeding 28 days after the first blm induction.

### Histopathology

Left lung lobes were formalin-fixed and paraffin-embedded (FFPE) to obtain longitudinal 5 µm thick Masson trichrome-stained sections for histopathological examination. The presence of lipid-laden alveolar macrophages (AMs) as pathological findings was evaluated in each group. The sections were scanned with NanoZoomer S60 (Hamamatsu Photonics K.K., Shizuoka, Japan) and imported into the Visiopharm Integrator System (VIS; version 2017.2.4.3387) for automated analysis of lung fibrosis using a VIS Analysis Protocol Package (APP). The sections were graded based on the Ashcroft scale (grades 0–8), as described by Ashcroft et al. (1988) [41] and modified by Hübner et al. (2008) [42].

### RNA extraction and sequencing

Four animals per group were randomly selected before sacrifice for transcriptomic analysis. Fresh right lungs of each animal were rinsed and immediately collected in RNAlater™ (Sigma, USA) for RNA preservation, and then they were homogenised in QIAzol® lysis reagent (QIAGEN, Netherlands) with gentleMACS™ Dissociator (Miltenyi Biotec, Germany). RNA was extracted from the lung homogenates using the QIAcube robotic workstation with the miRNeasy Mini Kit with an in-column DNase digestion protocol (QIAGEN, Netherlands). The RNA quality was measured using the Agilent 2100 Bioanalyzer system (Agilent Technologies, Santa Clara, CA, USA). The QuantSeq FWD (Lexogen) kit was used to prepare libraries, which were then sequenced on an Illumina NextSeq500 platform, producing at least 20 million reads/sample (75 bp Single End). The dataset is available with accession number GSE278200. Reads were aligned to the *Rattus norvegicus* genome (v. 6) using STAR [43], and counts were calculated using the HTSeq Python package [44] and Ensembl annotation (v. 101). ClustVis [45] was used to perform Principal Component Analysis (PCA) on the normalised read counts and the heatmaps were generated using the Morpheus software (developed by the Broad Institute, https://software.broadinstitute.org/morpheus).

### Transcriptomic data analysis

Differentially Expressed Genes (DEGs) were identified with the Limma-voom tool [46] using SAL as control group. Only genes with a minimum of 10 counts in at least 3 samples were considered. Genes were deemed as differentially expressed if the log_2_ fold-change (FC) was ≥ 1 or ≤ −1 and the adjusted p-value ≤ 0.05. Modules of co-expressed genes were identified using the “WeiGhted Correlation Network Analysis (WGCNA)” package in R [47]. Only protein coding genes with at least 10 counts in all the samples and 50 counts in 8 out of 12 samples were used for module construction. Modules were identified using the blockwise-modules function set at default, except for the following parameters: soft thresholding power = 9, networkType = “signed”, corType = “pearson”, TOMType = “signed”, minimum module size = 30, mergeCutHeight = 0.25, minKMEtoStay = 0.5. The correlation between gene expression module profiles and phenotypic traits (Ashcroft score, and % of Fibrosis) was evaluated using Pearson correlation. Metascape was used to identify significantly enriched pathways and processes in modules [48]. Enrichment of cell-type specific signatures within modules was assessed through gene set overrepresentation analysis. Specifically, an enrichment score was calculated that reflects the degree to which cell type marker genes are present more than would be expected (over-represented) among the genes assigned to a WGCNA module. The score was calculated by the ratio of the number of marker genes found in a WGCNA module divided by the total number of genes assigned to that module, to the total number of genes in the marker gene set divided by the total number of genes in the rat genome. The statistical significance of the enrichment score was estimated using Fisher’s exact test. We considered significantly enriched gene sets with an enrichment score ≥ 4 and a Fisher’s exact *p*-value ≤ 0.001. Cell-type specific markers were obtained from Mouse Cell Atlas [49–51], from Morse et al. [11], and from Hong et al [52]. Only genes with module membership ≥ 0.8 were retained in each module for this analysis. We used the same analysis to evaluate the enrichment of specific signatures from our previous work [8]. Specifically, we compared the module-specific signature of our previous time course investigation with our current study. Once again, only genes with module membership ≥ 0.8 were retained in each module for this analysis. Gene Set Enrichment Analysis (GSEA) [53] was employed to identify the enrichment for “nintedanib-sensitive” gene sets within the WGCNA modules. These gene sets originated from a series of relevant publications that analysed the gene expression effects of nintedanib in different types of ILD *in vitro* and *in vivo* models [10, 54–56].

### Assessment of lipid peroxidation by malondialdehyde (MDA)

Parallel experimental groups of animals were included for the MDA analysis on lung homogenates subdivided in SAL (n=6), BLM (n=9), and NINT (n=7) according to the procedure described above. Frozen right lobes were homogenized in 10 mL of ice-cold 1X PBS per g of tissue, with gentleMACS™ Dissociator. Lung homogenate was frozen/thawed three times and centrifuged (5000 g for 10 minutes) to obtain a supernatant cell lysate. The amount of MDA was determined with the Lipid Peroxidation Colorimetric Assay Kit (Abcam AB233471, UK) according to the manufacturer’s protocol.

### Graphs and statistical analysis

Venn-diagrams were performed using Venny 2.0 [57]. Histograms and statistical analysis were performed with Graphpad Prism v. 10.1.0 (GraphPad Software, San Diego, California, USA). The statistical analysis of histological examination and MDA measurement was conducted using the Kruskal–Wallis test, followed by Dunn’s multiple comparisons test.

